# Tissue-specific transfer-learning enables retasking of a general comprehensive model to a specific domain

**DOI:** 10.1101/2023.09.11.557208

**Authors:** Qing Li, Deshan Perera, Zhishan Chen, Wanqing Wen, Dinghao Wang, Jun Yan, Xiao-Ou Shu, Wei Zheng, Xingyi Guo, Quan Long

**Affiliations:** Department of Biochemistry & Molecular Biology, University of Calgary, Calgary, Canada; Department of Medicine & Biomedical Informatics, Vanderbilt University Medical Center, Nashville, USA; Department of Mathematics and Statistics, University of Calgary, Calgary, Canada; Physiology and Pharmacology, University of Calgary, Calgary, Alberta, Canada; Alberta Children’s Hospital Research Institute, University of Calgary, Calgary, Canada; Department of Medical Genetics, University of Calgary, Calgary, Canada; Hotchkiss Brain Institute, University of Calgary, Calgary, Canada

**Keywords:** Transfer learning, ChIP-seq, Breast cancer, Transcriptome-wide association study, Fine mapping

## Abstract

Machine learning (ML) has proven successful in biological data analysis. However, may require massive training data. To allow broader use of ML in the full spectrum of biology and medicine, including sample-sparse domains, re-directing established models to specific tasks by add-on training via a moderate sample may be promising. Transfer learning (TL), a technique migrating pre-trained models to new tasks, fits in this requirement. Here, by TL, we retasked Enformer, a comprehensive model trained by massive data, tailored to breast cancers using breast-specific data. Its performance has been validated through statistical accuracy of predictions, annotation of genetic variants, and mapping of variants associated with breast cancer. By allowing the flexibility of adding dedicated training data, our TL protocol unlocks future discovery within specific domains with moderate add-on samples by standing on the shoulders of giant models.

## INTRODUCTION

Machine learning (ML) including deep learning has been successfully applied to biological data analysis, especially multi-scale omics data integration^1-6^. However, to build effective ML models, one may usually have to carry out sophisticated training using a large sample. Such large samples may not be available in many domains, limiting the use of ML in broader spectrums. Moreover, large samples could contain heterogeneous components, leading to the final model (despite being extensively trained on massive data) being suboptimal for particular user cases. In recent years, in many fields, including natural language processing and image recognition, transfer learning (TL) has been widely used to migrate established (usually comprehensive) models to specific domains^7-13^. In general, a TL model is composed of three components: a base model (usually generally purposed) to start with, a sample (usually small) dedicated to a specific task, and a training procedure to re-train a subset of parameters of the base model towards the specific goal^14; 15^ (**Online Methods**). This whole procedure may be called “retask” to reflect the efforts of redirecting the original model to a specific task. Although the general procedure looks intuitive and straightforward, in practice, how to carry out such a retasking is an art relying on domain knowledge and trial-and-error experiments.

In genetics, genome-wide association studies (GWAS)^16-18^ provide preliminary links between genotype and phenotype. However, for the majority of GWAS-reported loci, potential causal variants and target tissues remain unknown. Towards this line, transcriptome-wide association studies (TWAS) improve GWAS by integrating gene expression data, leading to tissue-specific signals discovered in many diseases^19-21^.

Recent studies have shown that the power of association mapping analysis can be significantly improved by integrating informative priors learned from existing transcriptomic and epigenetic data^19^. In the same vein, we have shown that TWAS can be substantially improved if prior knowledge from specific biological mechanisms (such as transcription factors or TFs) is integrated^20-22^. However, these efforts are segmented based on different pieces of prior knowledge, and there is no routine procedure assisting the integration of most (if not all) existing omics-based prior annotations and, in the meantime, tailored to the specific disorder, not to mention the targeted specific tissue.

A significant step towards the above vision is the publication of Enformer, a highly sophisticated deep neural network model containing >200 million parameters, which digested massive epigenetic datasets, including a total of 5,313 tracks of TF binding, chromatin marks, and transcription profiles in various human tissues/cells^2^. By targeting the prediction of genetic variant effects on regulatory activities for each track of epigenetic profile, Enformer essentially provides integrated functional weights for any potential variants, which can be utilized by many downstream analyses^23; 24^. However, despite its comprehensive training data and highly sophisticated architecture, Enformer as a general-purpose model contains massive and heterogeneous information. As such, Enformer by itself may not be optimal for specific tasks in a target tissue.

In this work, we developed a protocol to carry out transfer learning (TL) to allow the addition of a small number of dedicated tracks to retask Enformer for specific tasks tailoring to desired domains. We selected breast cancer as an illustrating example. This is because of two rationales: First, Enformer includes only 35 tracks of epigenetic profiles in breast cancer-related tissues/cells. Second, disease-associated target tissue is lacking in Enformer’s training procedure. Third, alternation of TF-occupancy by cis-regulatory elements, a tissue-specific mechanism that has been shown important to many genes and cancers (including breast cancer)^20-22; 25; 26^.

In the breast cancer application, our TL framework to retask Enformer is achieved by further learning new 275 TF Chromatin Immunoprecipitation Sequencing (ChIP-seq) datasets in breast cells. We demonstrated that our TL outperforms Enformer in annotating target **t**issue-based **C**is-**R**egulatory **A**ctivities (tCRAs) for breast cancer. We showed that TL model outperforms Enformer in statistical predictions and functional annotations. Application-wise, we further showed that integrating breast related tCRAs significantly improves existing approaches in association mapping using TWAS in identifying independent risk signals for breast cancer. Thorough comparison to the base-model (Enformer) are also presented.

## RESULTS

### Outline of the Transfer Learning (TL) framework

A general framework of Transfer Learning (TL) re-trains a subset of nodes in a pre-trained model based on a small number of dedicated data (**Figure 1A**). Our TL is based on a pre-trained general-purpose model, Enformer, which is composed of four blocks: convolutional blocks, transformer blocks, pointwise convolution blocks, and output head layer (**Figure 1B**). We re-tasked Enformer to specificities of human transcription factor (TF) ChIP-seq data of 275 tracks (**Figure 1C**) by selectively updating a minimal set of parameters. Our rigorous experimentation and performance monitoring revealed that updating only a small subset of parameters—specifically, those in the final layer and the pointwise convolutional block—effectively leveraged pre-trained knowledge without unnecessarily complicating the model or risking overfitting. Similar to Enformer, the learned representations of genomic variants can be used for multiple purposes. In this work, a key application lies in estimating the influence of genetic variants on target track profiles, such as prediction of TF activities in the target tissue, e.g., breast, defined as target **t**issue-based **C**is-**R**egulatory **A**ctivities, or **tCRA**s (**Figure 1D**). We adopted standard training methods for deep learning models with Adam as the optimizer and Poisson negative log-likelihood loss function for the training of the TL model (**Supplementary Figure S1**). Detailed mathematical formulations of our TL framework, including the definition of tCRAs scores as well as their computational implementations and training procedures, are presented in **Online Methods**.

**Figure 1:**
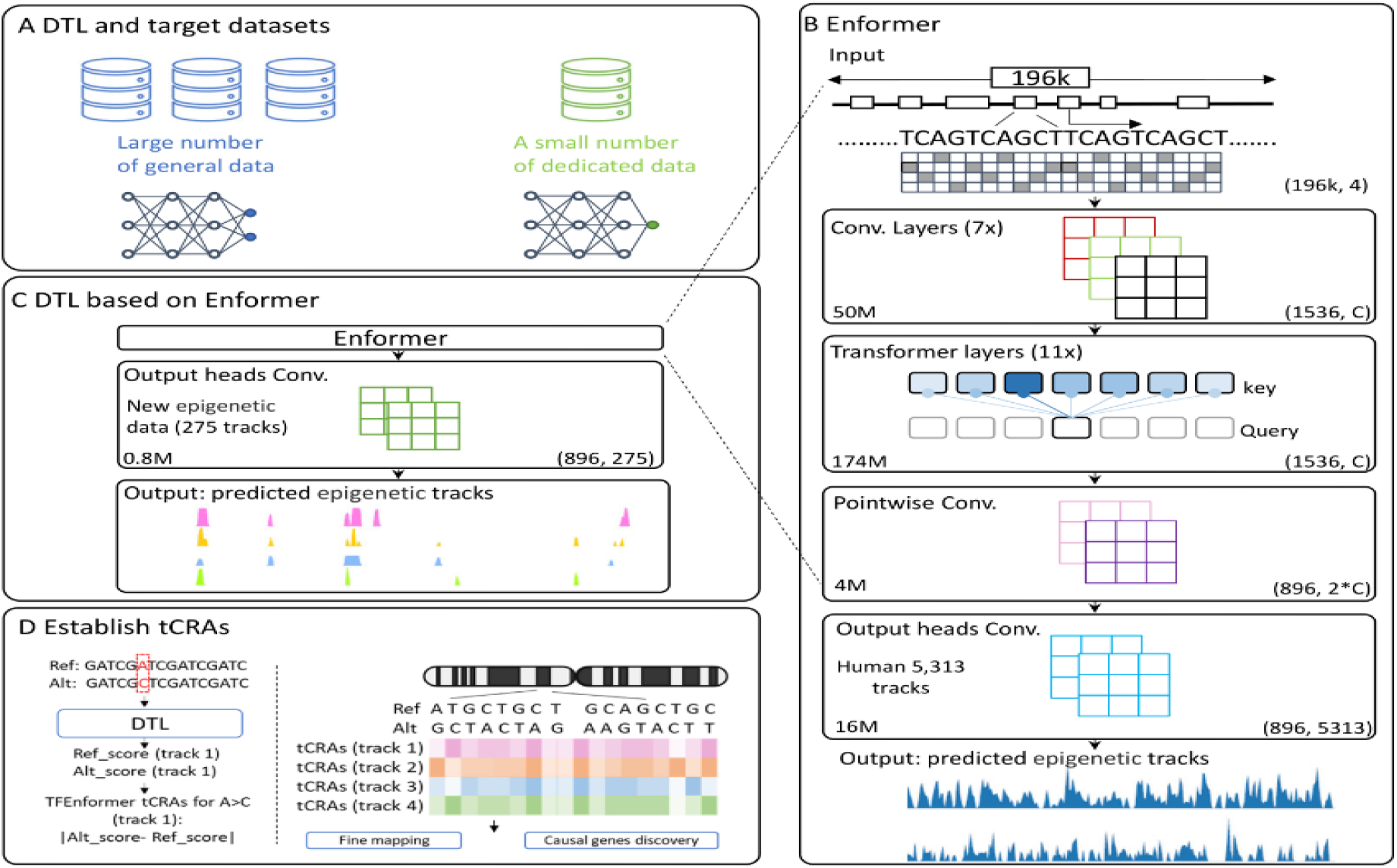
The workflow of our TL approach to establish tCRAs in breast cancer. **A)** A schematic of general TL: a small number of dedicated data may re-train a selected subset of nodes to retask an existing model. **B)** Enformer model was trained on input DNA sequences (196kb) to predict multiple human epigenomic tracks (5,313) at 128-bp resolution. The network is composed of four blocks (indicated by each rectangle) from convolutional layers, transformer layers, pointwise convolution, and output heads convolution. The output shapes of these blocks are given by tuples on the bottom-right corner, in which C (=1,536) indicates the number of channels in CNN. The number of trainable parameters for each block is listed on the bottom-left corner. **C)** Our TL used majority of existing Enformer architecture together with its trained parameters by retaining its input and first three blocks. The only layer undergoes re-training is the output heads convolution layers, tailed to target-tissue epigenetic datasets (i.e., 275 tracks of TFs ChIP-seq for breast cancer). **D**) Left panel: An illustration of tCRAs for a specific genetic variant estimated by calculating the differences of predicted regulatory activity value between reference allele (=A) and alternative allele (=C) of a variant. Right panel: based on TL outcomes, we generated an activity score for each genetic variant for each track, which can be utilized in downstream analyses including association study and fine-mapping.

### TL generated accurate statistical predictions

The accuracy of statistical predictions serves as the first metric in verifying whether the TL model indeed improves the model in the targeted domain, i.e., breast tissue. We first check whether our TL model can predict the TF ChIP-seq peaks, as exemplified in essential regions ESR1 and FOXA1 (**Figure 2A**). As an overall comparison between our re-tasked model and the original Enformer model, we compared the overall distribution of Person correlations (between predicted and the actual) for all tracks for 15 TFs shared by Enformer and TL on the hold-out validation dataset (**Online Methods**), showing a significantly higher distribution of the TL model than Enformer (**Figure 2C**). To further characterize the contribution of each TF to the above superior overall distribution, by stratifying the tracks for different TF, the individual average correlations of all TFs are depicted (**Figure 2B**), ranging from around 0.34 (ESR1) to 0.75 (E2F1). A head-to-head comparison between TL and Enformer shows that, except for four TFs, the mean correlation of TL consistently outperforms its counterpart in Enformer (**Figure 2D**). Evidently, this initial check of statistical accuracy in predicting ChIP-seq signals illustrates that the TL model, trained on regulatory data from 275 TF ChIP-seq tracks, successfully improved the specificity towards the transferred domain.

**Figure 2:**
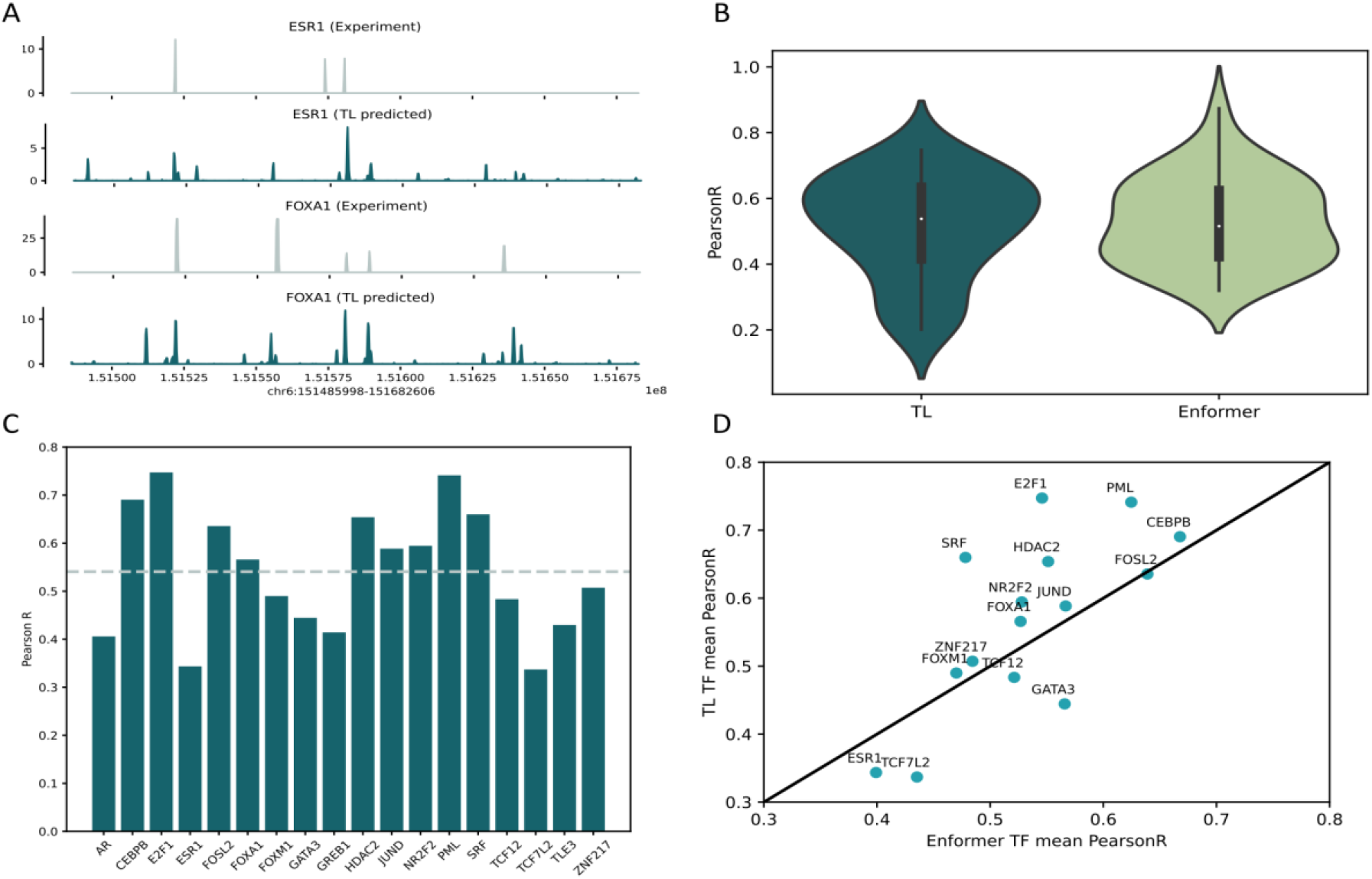
Performance of TL in statistical predictions. **A)** Experimental and TL Predicted CHIP-seq Track Profiles for ESR1 and FOXA1. The genomic region Chr6:151485998-151682606 from the loci chr6q25.1 was selected for analysis. This region encompasses the SNP rs2046210 (chr6:151627231), and previous research^27^ reported ESR1 to be the TF located within a 1M proximity of this SNV. **B)** Bar plot to show the average Pearson correlation coefficients (by the number of tracks) between predicted outputs and true outputs for each TF. The grey dash line is the average Pearson R across TFs. **C)** Violin plots to show Pearson correlation coefficients between predicted outputs and true outputs for all tracks shared by TL and Enformer. **D)** Scatter plots to show the averaged Pearson correlation coefficients (by the number of tracks) for TFs shared by TL and Enformer.

### SNVs prioritized by TL-derived tCRAs are enriched in multiple functional annotations

The first application of TL is to assess the impact of genetic variants on target track profiles, a metric we termed tCRAs (**Online Methods**). Mathematically, the calculation of tCRAs is to reflect the difference in predictive outcomes by switching the reference allele to an alternative allele using a bin centralized by the focal variant; and it is dichotomized into 1 or 0 by comparing to the population mean (**Online Methods**). In this study, to verify the TL model from many aspects, we used data from Breast Cancer Association Consortium (BCAC), a large international consortium studying genetic basis of breast cancer containing 123K cases and 106K controls^28^ (**Online Methods**). Here we applied both TL and Enformer to evaluate the tCRAs (designated as 0 or 1) for 9.9 million SNVs that passed quality control in the BCAC GWAS summary statistics. We then prioritize SNVs using their number of tracks with tCRAs being 1, guided by the intuition that SNVs with more tracks with tCRA=1 play more critical roles than those with fewer tracks with tCRAs=1 (**Online Methods**). As the number of tracks for TL and Enformer is different, we, therefore, adjusted the cutoffs of a number of tracks separately for TL- or Enformer-annotated to ensure a roughly equal number of SNVs were in corresponding prioritized variants subsets. This process resulted in five subsets of SNVs: 500K, 1M, 1.5M, 2M, and 3.5M, with a detailed breakdown of SNVs provided in **Supplementary Table S1**.

A fundamental hypothesis driving the development of the TL model is that the TL may improve the performance in a specific disease and mechanism compared to the general-purposed Enformer model. We analyzed the enrichment from three aspects of functional annotations, i.e., functional region, association to a disease, and cell-type specific (**Online Methods**), reaching observations that substantiate this assumption (**Figure 3**). First, we looked at whether the prioritized SNVs are located in breast cell line-specific enhancer regions, and we observed that all subsets showed that TL-derived prioritizations enjoy higher enrichment (**Figure 3A**). Second, we checked the overlap between the BCAC-reported SNVs associated with breast cancer and the prioritized SNVs subsets and again detected that TL tCRAs prioritized subsets captured more GWAS risk SNVs than Enformer (**Figure 3B**). Third, we checked the percentage of enhancer-located SNVs by stratifying the cell types and found that the distribution in breast-related cells is higher in non-breast cells (**Figure 3C**). Although this pattern is also observed in Enformer tCRAs prioritized SNVs subsets, the design is quite subtle (**Figure 3D**) and not as propounded as the TL case. Taking together, we demonstrated the capacity of TL to retask the comprehensive Enformer model to breast tissue with a focus on TF binding, suggesting its utility in studying the genetic foundation underlying disease etiology.

**Figure 3:**
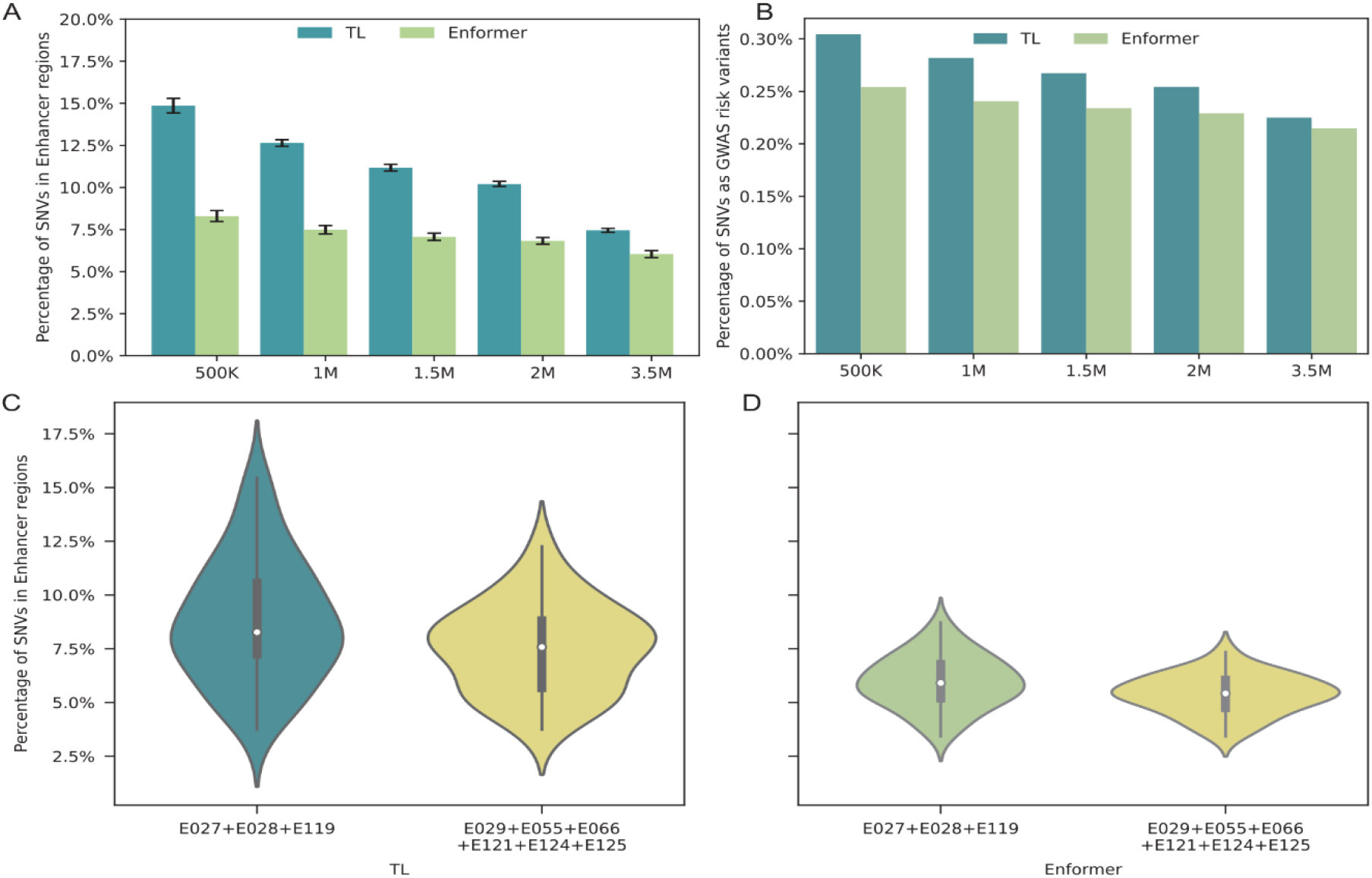
Functional enrichment of SNVs in top tCRAs. **A)** The proportion of SNVs located within enhancer regions for the five TL- or Enformer-prioritized SNV subsets, using their corresponding tCRA scores. Intervals were calculated by three breast cell lines. **B)** The proportions of TL- or Enformer-prioritized SNV subsets that are overlapped with of BCAC GWAS risk SNVs (with P-values < 5e-8). **C)** Violin plots for proportion of SNVs located in enhancer regions of all TL-prioritized tCRAs within breast-specific cell lines (E027+E028+E119) and other cell lines (E029+E055+E066+E121+E124+E125). **D)** The same as C but using Enformer-prioritized tCRAs.

### TL-prioritized variants improve genotype-disease association mapping in TWAS

As a model of integrating gene expression data in GWAS, TWAS analyzes the correlation between disease phenotype and an aggregation of cis-genetic variants selected by gene expressions^29; 30^. A conventional TWAS protocol usually uses a regularized multiple regression (e.g., elastic net) to select SNVs and their weights in the form of training a gene expression prediction model (in a reference dataset such as GTEx^31^; and then apply this model to “impute” the expression in the GWAS dataset for assessing the association (**Online Methods**). Previously we have demonstrated that the pre-selection of variants based on functional priori^20; 21^ for the predictive model training substantially improve the power of TWAS. In this study, we leveraged five SNVs subsets prioritized by tCRAs derived by TL and Enformer respectively to learn if machine-learned functional scores bring further improvement on the breast cancer data generated by BCAC (**Online Methods**).

We first generated the baseline using the default Fusion method, i.e., all the 9.9M SNVs from BCAC without any prioritization are supplied to the analysis and the genome-wide significant genes after Bonferroni correction (**Online Methods**) are reported. This result, depicted as the dashed line in **Figure 4A** acted as a benchmark for the comparisons. Interestingly, SNVs subsets prioritized by Enformer-derived tCRAs do not improve the model overall: it produced slightly higher numbers in the 1.5M, 2M and 3.5M subsets and even lower numbers in 500K and 1M subsets. This is consistent to our hypothesis that a general-purposed model might not be optimal for dedicated tasks. In contrast, TL re-tasked model shows substantial improvement by discovering higher numbers of genes over the baseline and also Enformer cross the board.

**Figure 4:**
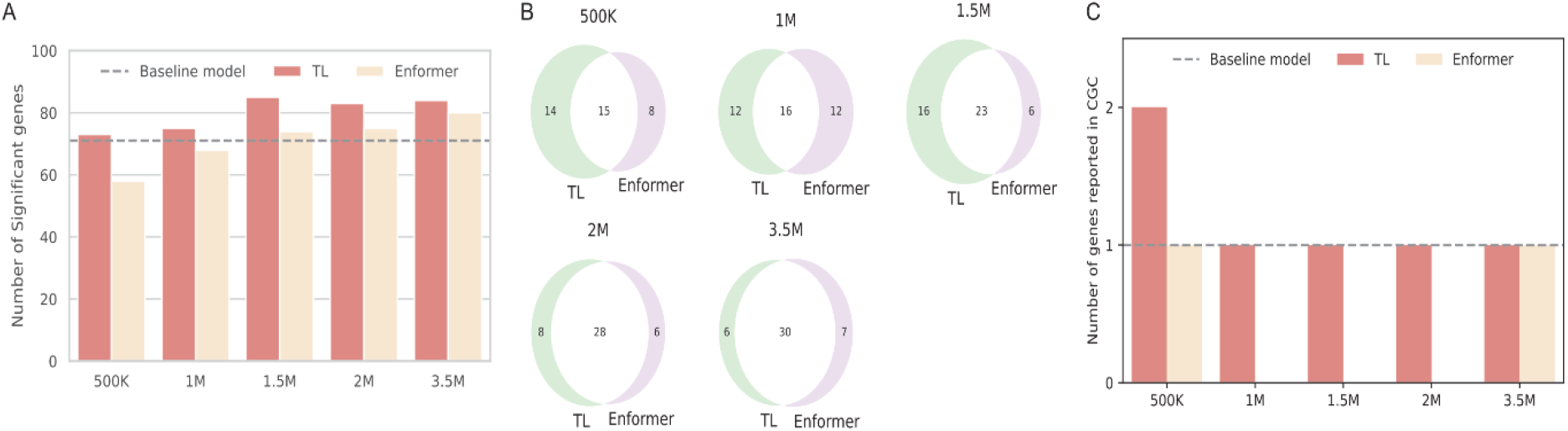
TL-derived tCRAs improved TWAS results in breast cancer. **A)** Numbers of Bonferroni-corrected significant genes identified by TWAS using TL- or Enformer-prioritized five subsets of SNVs. The dashed line indicates the performance of base model that do not use any functional prioritizations. **B)** Numbers of significant genes validated by the DisGeNET database across five TL- and Enformer-prioritized SNV subsets. C) Numbers of significant genes validated by the CGC database across five TL- and Enformer-prioritized SNV subsets.

Next, we verify the relevance of these discovery using reported genes in DisGeNET^32^, a comprehensive disease-gene database, and Cancer Gene Census^33^, a dedicated database for cancers. Our investigation revealed that the five TL subsets (500K, 1M, 1.5M, 2M, and 3.5M) - based analysis contained 29, 28, 39, 36, and 36 genes reported in DisGeNET, respectively, whereas the Enformer-based analysis contained only 23, 28, 29, 34, and 37 genes, respectively (**Figure 4B**). Of note, the baseline model contained 32 genes. The CGC reported far fewer genes compared to DisGeNET (42 vs. 7,083), and consequently, the number of significant CGC-reported genes in the five subsets is also smaller. However, the same trend is pronounced (**Figure 4C**): In the TL-based analysis, the 500K subset-based analysis has 2 genes verified by CGC, while each of the remaining four TL subsets-based has 1 gene verified by CGC. In contrast, for the Enformer-based analysis, only 500K and 3.5M-based has 1 gene verified by CGS with all the other three zero. The baseline model has 1 CGC-verified gene. These collectively underscore that TL tCRAs enhance the ability to identify disease susceptibility genes specific to certain diseases, as illustrated by our study on breast cancer.

## DISCUSSION

In biological research, although TL has been applied to some application such as single-cell analysis, systematic retasking of comprehensive models based on multi-scale omics data has not been carried out. This work represents the first study that proposes a framework to retask a comprehensive model. Using breast cancer as an illustrating example, we have thoroughly tested the transferred model in contrast to the base model Enformer regarding the statistical predictions, annotation of variants, and downstream association mapping. An immediate future work may be testing this TL framework on other diseases.

It was reported that functional annotation based on epigenetic data from non-target tissues could introduce a wide range of cellular functions for nonspecific elements not intrinsic to the disease state^34^. As such, training a multi-scale data integration model will frequently suffer from the trade-off between too heterogenous samples and too narrowed scope. Here the development of our TL framework split the task into two steps. First, one can train a comprehensive model using whatever data are available to integrate all sources of information. Then, the researchers with dedicated data and expertise can retask to general model using our framework to apply the integrated scores to their specific applications.

The applications of transferred model are not limited in TWAS demonstrated in this work. All the applications that require interpretation of variants may benefit from this transferred model. for instance, fine-mapping is a promising field that integration of priori knowledge-based scores may play an important role^19; 23^. Additionally, such scores may be used to train more accurate polygenic risk scores^35; 36^ for clinical use.

Although our TL framework provides a computational protocol to retask Enformer, the performance of transferred models still need manual inspections and fine-tuning to optimize. An interesting future work might be integrating this framework with theoretical research on the transferability^37^ of existing models to quantify the feasibility of such a transfer learning and automate some of the parameter tuning steps.

## ONLINE METHODS

### Data sources

#### The breast transcription factor (TF) chromatin immunoprecipitation and sequencing (ChIP-seq) data

were obtained from the Cistrome database^38^. In total, there are 275 tracks covering 18 TFs (AR, CEBPB, E2F1, ESR1, FOSL2, FOXA1, FOXM1, GATA3, GREB1, HDAC2, JUND, NR2F2, PML, SRF, TCF12, TCF7L2, TLE3, ZNF217) (**Supplementary Table S2**).

Adhering to the quality control (QC) guidelines from the Cistrome database, we ensured the inclusion of high-quality TF ChIP-seq datasets for subsequent analyses. The sample’s QC standards comprised several indicators: a FastQC score, representing the median sequence quality, had to exceed 25; the UniquelyMappedRatio, indicative of the mapping quality, had to surpass 0.6; the PBC value, employed to detect potential PCR over-amplification, had to exceed 0.8; the PeaksFoldChangeAbove10, denoting the number of peaks identified by MACS2 with a fold change of 10 or more, had to exceed 500; the FRiP score, representing the fraction of non-mitochondrial reads in the peak region, had to be greater than 0.01; and, finally, the PeaksUnionDHSRatio, utilized for assessing the annotation level QC, had to be greater than 0.7.

#### The Genome-Wide Association Study (GWAS) data

for breast cancer were downloaded from the Breast Cancer Association Consortium (BCAC), an international initiative dedicated to identifying genetic risk factors associated with breast cancer^28^. This dataset encompasses GWAS data for 122,977 cases and 105,974 controls, all European descent. SNVs with minor allele frequency > 0.01 (number of qualified SNVs is 9,898,466) are utilized for analyses in this study.

#### The GTEx expression reference dataset

We downloaded gene expression and individual genotype data from the GTEx portal^31^. We processed the normalized gene expression data using the Probabilistic Estimation of Expression Residuals (PEER) analysis^39^, which adjusts for potential confounding variables. We used 30 PEER factors for our downstream model building based on the GTEx recommendation for the breast tissue. We also QC-ed the GTEx genotype dataset using PLINK software^40^, removing variants with high missingness (remove SNVs with less than 10% genotyping rate), low minor allele frequency (MAF < 0.01), and notable deviations from Hardy-Weinberg equilibrium (HWE P-value < 1 × 10^−6^). There are 9,527,843 SNVs remained after filtrations.

#### Functional enrichment annotation datasets

To investigate the functional role played by subsets of SNVs, we downloaded genomics regions with annotated by 15 chromatin states in Roadmap Epigenomics project^41^. They used ChromHMM v1.10^42^ to discern 15 chromatin states, available as BED files on their website. To verify the evidence of cancer-related susceptibility genes, we used established databases including DisGeNET^32^ and Cancer Gene Census (CGC)^33^ from the Catalogue of Somatic Mutations in Cancer (COSMIC) website.

### The transfer learning (TL) algorithm

#### A general framework of transfer-learning

(TL) starts from a pretrained model *M* = (*T, D*, Θ) where *T* is the topology of the (usually deep) network, *D* is the (usually massive) datasets, and Θ is the parameter vector trained by a loss function. A TL *M*^*TL*^ = (*T*′, *D*′, Θ_0_, Θ′) will adjust some of the nodes from *T*, leading to the revised network *T*′; and supply additional data (*D*′) to re-train a subset of the parameters Θ′, retaining the rest parameters Θ_0_ = Θ − Θ′. Note that the number of parameters retained in *M*^*TL*^, i.e., |Θ′| indicates how authentic the new model will be consistent to the original model, in contrast to the level of dedication to the new task.

#### A brief outline of the original Enformer model

Enformer^2^ is a profoundly advanced deep neural network with self-attention mechanism, encompassing Θ = over 200 million parameters. The training data of Enformer includes both humans and mouse. Here we briefly introduce the human part. The human training data *D* = (*D*_*input*_, *D*_*outpuit*_), where *D*_*input*_ are organized by dividing human genome into bins, each of which contains a small segment of 196,608 base pairs (bp) from the HG38 human reference genome. This organization leads to 38,171 bins, distributed as 34,021 for training, 2,213 for validation, and 1,937 for test. To expedite mathematical operations in Enformer, the one-hot-encode DNA sequences (A = [1,0,0,0], C = [0,1,0,0], G = [0,0,1,0], T = [0,0,0,1], N = [0,0,0,0]) is adapted, resulting in a 196,608 × 4 input matrix. At the other end, *D*_*outpuit*_ = 5,313 genomic tracks for humans as outputs to be predicted. The data structure of *D*_*outpuit*_ is a matrix with dimensions 896 × 5,313, where 5,313 represents the total number of genomic tracks associated with transcription factor (TF) binding, chromatin marks, and transcription profiles across various human tissues and cells; and 896 = (196,608-40,960*2)/128) is the result of deducting twice the value of 40,960 (the base pairs removed from both ends of the sequence) from the length of the input DNA bins (196,608), and then dividing by 128, indicating that each of the 896 values represents an averaged profile over 128 DNA base pairs. The architecture of Enformer, *T* is strategically divided into three core components: *T*_1_: convolutional blocks with pooling, *T*_2_: transformer blocks, and *T*_3_: a cropping layer succeeded by pointwise convolutions associated with *T*_4_: organism-specific network heads. Through the training of Enformer, the parameters are refined by minimizing the Poisson negative log-likelihood loss function, which assesses the discrepancy between the matrix predicted by Enformer and the actual values in the target matrix.

#### Our TL algorithm and additional data

The design of our TL model aligns closely with the Enformer architecture, incorporating its three main components. Our adjustment is described below: first our training sample *D*′_*output*_ = 275 breast TF ChIP-Seq data described in “Data sources” (**Supplementary Table S2**). The *D*_*input*_ is the human genome part of Enformer (which originally contains mouse as well). The network topology *T* is only modified at *T*_4_ by eliminating the two output heads of Enformer, which predict organism-specific tracks, and replacing them with a new output head tailored to our 275 target tracks. We also added a one-dimensional convolutional layer with 275 filters (each with width =1), resulting in 275 output channels, each corresponding to a distinct track. This modification is crucial because each filter, or “kernel,” a small matrix employed in extracting specific features from input data via convolution, generates a single channel. These channels represent different feature maps or filters that capture unique patterns or features in the input data. Mathematically, a one-dimensional convolutional layer with a kernel size of 1 is equivalent to a fully connected layer. So in reality, this 1D convolutional layer function is realized by the linear layer function (snt.Linear()). Furthermore, we applied the ‘softplus’ function, similar to Enformer, as an activation function. In total |Θ′|, the number of adjusted parameters is approximately 20∼30M.

#### Re-training procedure of the TL model

We retained parameters from Enformer to TL for convolutional blocks, transformer blocks, and pointwise convolution blocks and randomly initialized parameters in the output head layer. Mirroring Enformer, the Poisson negative log-likelihood loss function was employed to learn regulatory signals from the TF occupancy in ChIP-seq and to predict the effects of genetic variants on alterations in TF ChIP-seq activities. The entire TL analysis was executed using Sonnet v2 and TensorFlow (v2.4.0), and training was conducted on a single GPU card (A100) with a batch size of 64 for a limited number of steps. Additionally, due to memory limitation, gradient accumulation^43^ was adapted, allowing the adjustment of parameters across multiple batches to be aggregated. The Adam optimizer from Sonnet v2 was employed, maintaining a learning rate of 0.0001, consistent with the fine-tuning step in the Enformer model.

Within our TL, the fixed parameters, Θ_0_ contains parameters from the majority of layers, while fine-tuning parameters are in only a few layers near the output to accommodate new tasks. In this study, we meticulously explored the number of layers to be fine-tuned during the transfer learning process. To do this, we conducted experiments with various configurations, updating parameters in different numbers of layers in the TL model, and then assessed the model’s performance using validation datasets. As depicted in **Supplementary Figure S2**, the TL architecture, from output to input, encompasses one linear connected layer, a pointwise convolutional block, 11 transformer blocks, and additional miscellaneous blocks. Therefore, we began the transfer learning procedure from the linear layer, the one nearest to the output, with the goal of determining the optimal configuration that optimally balances model performance. We conducted four tests, incrementally increasing the number of layers to be fine-tuned: 1) updating parameters only in the final linear fully connected layer; 2) updating the final layer and the pointwise convolutional block; 3) updating the final layer, the pointwise convolutional block, and one transformer block; 4) updating the final layer, the pointwise convolutional block, and three transformer blocks. The optimal model in transfer learning was selected based on two metrics estimated in the validation datasets: 1) Pearson correlation scores computed for 5,313 tracks, and 2) Loss calculated between predicted and true values.

### Construction of target tissue-based Cis-Regulatory Activities (tCRAs)

#### TL-derived tCRAs

Based on the re-tasked training above, we define tCRAs (**Figure 1D**) to reflect the impact of genetic mutations on the cis-regulatory activities below:

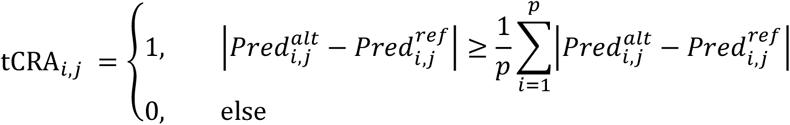

where, tCRA_*i,j*_ is the tCRA for *i*-th SNV, *j*-th track. 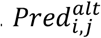 is the predicted value of the *j*-th track when the *i*-th SNV has the proposed alternative allele, which is in the context of a bin of DNA sequence of 196,608bp; 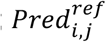 is the predicted value of *j*-th track when the *i*-th SNV has the reference allele; *p* is the total number of SNVs utilized to estimate effect sizes.

Specifically, for a single track, TL produces an output vector consisting of 896 values. The reference/alternative allele is located just one position to the right of the DNA sequence’s center, which means that its corresponding output value 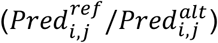 is also centered in the output vector. The effect of *i*-th SNV (ref -> alt) on *j*-th track is estimated by absolute value of discrepancy between 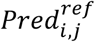 from 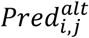. We then dichotomized this discrepancy based on the average values of all SNVs effects. For each distinct track and SNV, a tCRA value of 1 is designated if SNV’s absolute effect size surpasses the averaged effect size of the track; otherwise, the tCRA is denoted as 0. We use this dichotomized value instead of simply using the value of 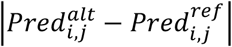 because of the following reasons: (1) This process of dichotomization harmonizes the tCRAs with other standard functional annotations, such as those from PolyFun^19^, facilitating the integration of tCRAs to enhance the prior causal probability in fine-mapping. Moreover, the dichotomized tCRAs simplify the calculation of accumulated effects of multiple tracks by counting the 1s in all tracks.

#### Enformer-derived tCRAs

To compare TL to Enformer, we also defined the Enformer version of tCRAs. Specifically, we downloaded the pre-calculated variant effects from the Enformer GitHub. Using the Enformer predictions of 12 TF tracks of the breast tissue (**Supplementary Table S3**), we defined the Enformer analogue of tCRAs based on the same formula above.

#### Prioritization of SNVs using tCRAs

In this work, based on the levels of strength of tCRAs, we prioritized five subsets of SNVs (500K, 1M, 1.5M, 2M and 3.5M). The dichotomization mentioned above makes the definition of the cutoffs more straightforward. We rank all SNVs based on the number of tracks in which the tCRA is defined as 1. Then all SNVs are ordered by their counts of tCRAs being 1. Note that the orders or SNVs are different if using TL- or Enformer-derived tCRAs. By tuning the cutoff of number of tracks in two different models, TL and Enformer, we ensured approximately the same number of variants into the downstream analysis, e.g., enrichment. The detailed cutoffs (number of tracks) and exact number of SNVs prioritized in each set are presented in **Supplementary Table S1**).

### Assessment of the statistical performance of TL

#### Initial evaluation using overall predictive accuracy

We evaluate the performance of TL model by calculating Pearson R for 1,937 DNA sequences from the test dataset. For a given DNA sequence with length of 196,608 units, TL yields an output matrix of 896 × 275 values. Here, the 896 values are predicted profiles, with each individual value in the vector corresponding to an averaged profile across a 128bp bin. The test datasets contained 1,937 bins, each 196,608 units long. These were consolidated into a stacked matrix of 1,737,552 × 275, where 1,737,552 is the product of 1,937 and 896. For each of 275 tracks, a Pearson R correlation is calculated for between a vector of length 1,737,552 with true values and a vector of length 1,737,552 with predicted values from the TL model. We then evaluated Pearson R correlation for each TF based in the TL model. Each of 1,937 DNA sequences contains 196,608 units in the test dataset. Each track yields one Pearson’s R value, leading to a total of 275 Pearson’s R values. For each TF in the TL model, the mean values are computed, taking into account the number of tracks associated with that TF in **Figure 2B**.

#### Identifying tracks that are comparable between TL and Enformer models

The TL model incorporates 275 ChIP-seq tracks, which are distributed across 18 distinct transcription factors (TFs). To ensure a compatible analysis, we meticulously selected 96 tracks from Enformer that aligned with 15 of the 18 TFs from TL, as detailed in **Supplementary Table S4**. It is important to note that these 96 tracks were not originally derived from breast tissue. Nevertheless, previous research suggests that the variations in transcriptional activity of TFs across different tissues are subtle^44^, thereby justifying our assumption that the tracks are sufficiently comparable for our analysis. Notably, the quantity of tracks associated with each TF varied between Enformer and TL. To address this, we adopted a conservative approach by selecting the smaller number of tracks available for each TF from either Enformer or TL, resulting in a final selection of 40 tracks. A detailed breakdown of the final track counts for each TF from both Enformer and TL is as follows: FOXA1 (11), ESR1 (7), CEBPB (4), FOSL2 and GATA3 (3 each), FOXM1 and NR2F2 (2 each), HDAC2, JUND, PML, SRF, TCF12, TCF7L2, ZNF217 and E2F1 (1 each).

#### Comparison of statistical predictions using TL/Enformer comparable tracks

These above 40 selected tracks correspond to 15 transcription factors (TFs), thereby resulting in a refined matrix of 896 × 40 values. Our test datasets included 1,937 DNA sequences, each encompassing 196,608 units. These sequences were amalgamated into a comprehensive stacked matrix of 1,737,552 × 40, where 1,737,552 is the result of multiplying 1,937 by 896.

Following this, for each of the 40 tracks, we calculated the Pearson’s R correlation between the values predicted by the model and the actual values obtained experimentally. This exercise resulted in a set of 40 Pearson’s R values for both the Enformer and the TL models. These 40 Pearson’s R values are illustrated in a violin plot (**Figure 2C**) as well as a QQ plot (**Figure 2D**), which is based on the average values of Pearson’s R for each TF.

#### Comparison of functional relevance of subsets of SNVs

Functional enrichment analyses assist in evaluating significant overlap between the SNVs in our selected sets and biological functional regions, which may imply a functional role of these SNVs sets in disease etiology. Typically, enrichment analysis was conducted by calculating the percentage of functional enrichment based on the ratio of functional SNVs to the total SNVs in each subset. We considered a genetic variant as a functional SNV if it resided within the functional regions (e.g., enhancer regions). In this study, we engaged three cell lines from breast tissue and six randomly selected cell lines from other tissues (**Supplementary Table S5**). Each cell line presents 15 chromatin states, but we primarily focused on the seventh chromatin state, which corresponds to the enhancer regions as determined by the NIH Roadmap. We collected all genomic regions spanning the enhancer regions, then computing both the count and percentage of SNVs from the five SNV subsets prioritized by their contributions from tCRA (described in the paragraph of **Prioritization of SNVs using tCRAs**.)

### Transcriptome-wide association study (TWAS) analysis

#### TWAS protocol

We used Fusion^30^ to carry out TWAS analysis, which is a 2-step protocol plus a permutation test. We outline the Fusion procedure below.

In the first step, we trained a predictive model using GTEx expression and genotypes. Particularly, the cis-variants (500Kb flanking region) are used as input terms and the elastic-net is used as the predictive model:

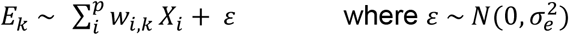

where *E*_*k*_ a vector representing the gene expression of the *k-th* gene; *X*_*i*_ is a vector illustrating the genotype of the *k*-th variant (*X*_*i*_=0, 1, or 2) and *p* represent the total number of variants in the 500Kb flanking region for the *k*-th gene; *w*_*i,k*_ is the effect size of *i-th* variant for *k-th* gene, and ɛ is the residual with 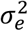 as its variance. The elastic-net penalty, a mixture of *L* _1_and *L*_2_ regularizers is used optimize (*w*_*i,k*_) for SNVs.

In the second step, the above trained weights 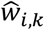 will be applied to the summary statistics of a GWAS data (in this project, BCAC data):

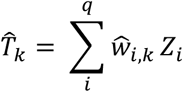

where *Z*_*i*_ is standardized effect size of *i-th* genetic variant for a trait at a given *cis* locus from GWAS summary statistic files, *q* represents the total number of variants in cis locus of the *k-th* gene in target dataset, and 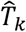 is the estimated Z-score (reflecting the association) of the expression and trait in the target dataset.

The significant of 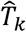 can be further estimated through permutation. In null hypothesis of no SNP-trait association 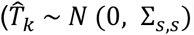, where Σ_*s,s*_ is the covariance among all SNPs (LD)) and 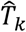 is calculated as *WZ* /(*W*Σ_*S,S*_*W*^*t*^)^1/2^. However, under an alternative model where 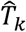 is drawn from a non-zero mean distribution, the distribution of 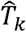 depends on both Z-scores of SNVs for a trait and *W* weights of SNVs for gene expression. To account for this, a permutation test is conducted, where expression labels are randomly shuffled and the 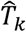 is computed against this permuted null distribution. This tests for an expression-trait association conditional on the observed GWAS statistics. The test is run 1,000 times for each gene in the TWAS analysis to compute a P value against this empirical null distribution.

#### Our TWAS analysis using prioritized SNVs subsets

In this study, we altered the above conventional TWAS protocol by supplying the elastic-net training with only prioritized SNVs. The GTEx and BCAC GWAS summary statistics data mentioned above is used for this TWAS analysis. The five subsets prioritized by TL-derived tCRAs are used for the assessment. As a comparison, Enformer-derived tCRAs are also tested. We also tested the baseline model without any prioritization. In all analysis, the genes with predictive R^2^ > 0.01 are retained for the association mapping and the genes with Bonferroni-corrected P-values lower than 0.05 are reported as significant.

## Data and code availability

TL is publicly available in our GitHub: https://github.com/theLongLab/Transfer-Learning BCAC: http://bcac.ccge.medschl.cam.ac.uk/

Cistrome TF ChIP-seq database: http://cistrome.org/Roadmap project 15 chromatin states: https://egg2.wustl.edu/roadmap/web_portal/chr_state_learning.html#core_15state

Bed files for nine cell lines from Roadmap project: https://egg2.wustl.edu/roadmap/data/byFileType/chromhmmSegmentations/ChmmModels/coreMarks/jointModel/final

COSMIC Cancer Gene Census: https://cancer.sanger.ac.uk/census

DisGeNet: https://www.disgenet.org/search, <u>Concept Unique Identifier (CUI) used for breast cancer are C0678222 and C0006142.</u>

Enformer: https://github.com/deepmind/deepmind-research/tree/master/enformer

## Author contributions

(Q.L.= Qing Li and Q.L.2 = Quan Long) Q.L., X.G., Q.L.2; Developed the software: Q.L.; Analyzed data: Q.L., D.P., Z.C., W.W., D.W., X.G., Q.L.2; Provided data, protocols and comments: J.Y. X.-O.S. W.Z.; Wrote the manuscript: Q.L, X.G, Q.L.2 with contributions from all co-authors.

## Supplementary information

**Supplementary Figure S1:**
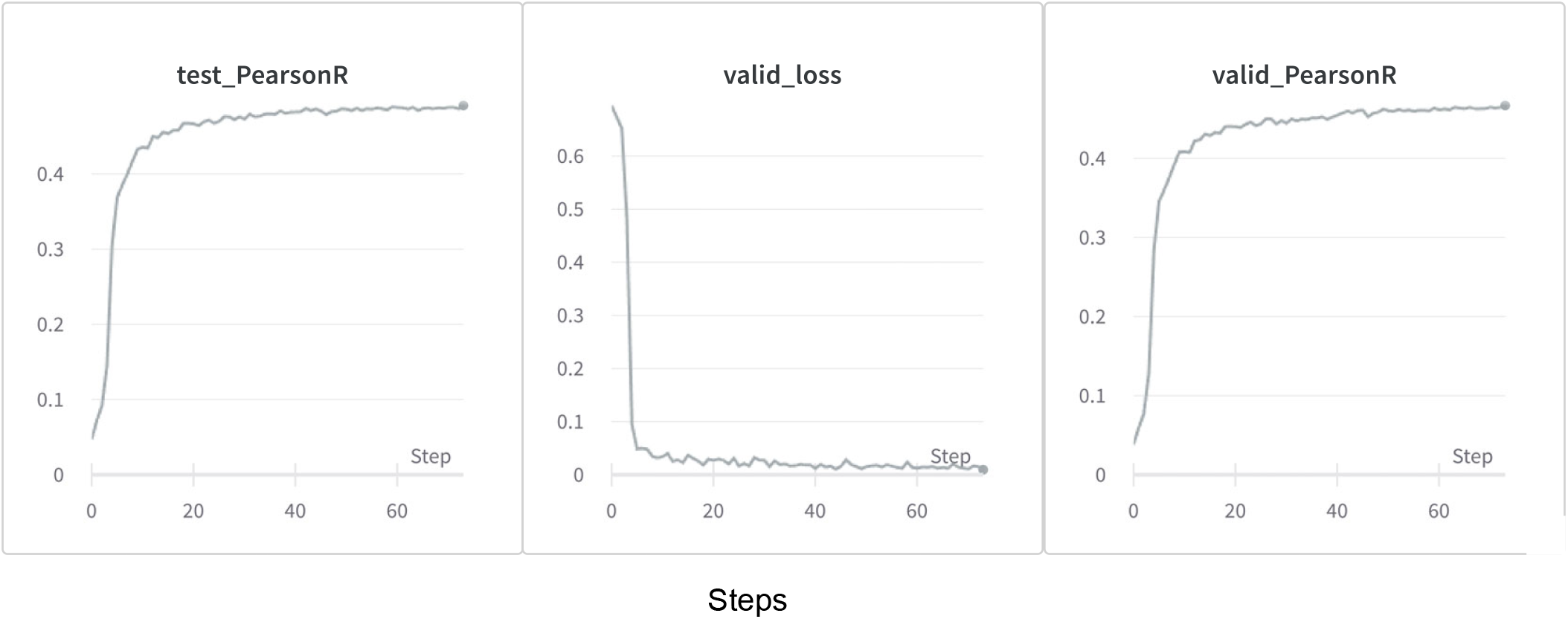
Transfer learning metrics. Pearson correlation in (A) test and (B) validation datasets and (C) the loss function in validation datasets.

**Supplementary Figure S2:**
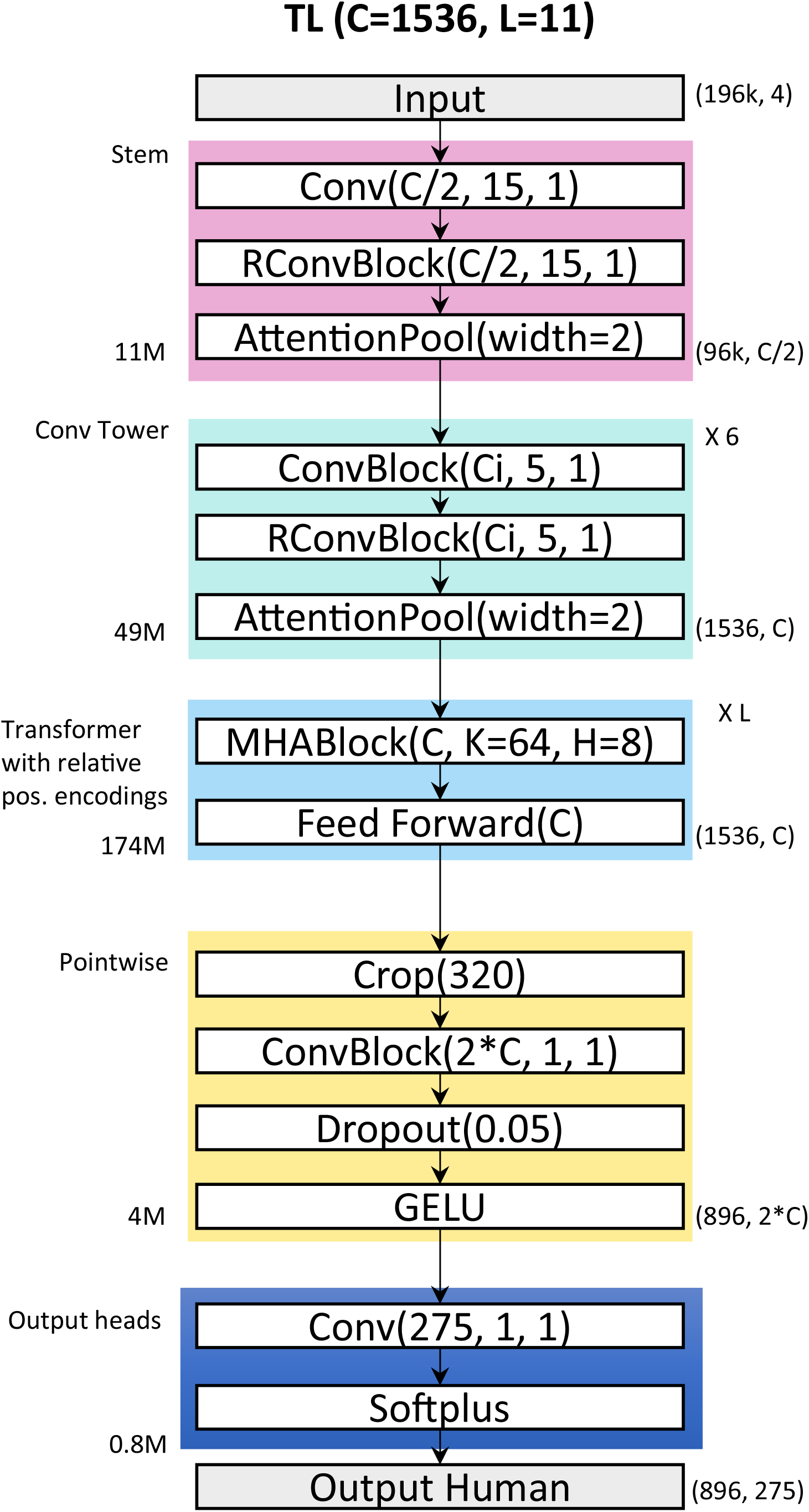
TL model architecture. The TL model is segmented into five distinct blocks, each composed of multiple layers. The output shapes, excluding the batch dimensions, are denoted by tuples situated to the right of each block. ‘L’ stands for the number of transformers and is equivalent to 11, while ‘C’ denotes the number of channels, amounting to 1,536. The TL model is characterized by a single output head, fine-tuned to cater to its designated TF tracks.

**Supplementary Table S1: Detailed cutoffs (number of tracks) and exact number of SNVs for five subsets from TL and Enformer**

**Supplementary Table S2: Breast 275 TF tracks from Cistrom (Transfer Learning)**

**Supplementary Table S3: Breast 12 ChIP-seq TF tracks (Enformer)**

**Supplementary Table S4: 96 tracks from Enformer that aligned with TL**

**Supplementary Table S5: 9 cell lines selected from the Roadmap project**

## Reference

1. Zhou, J., and Troyanskaya, O.G. (2015). Predicting effects of noncoding variants with deep learning-based sequence model. Nat Methods 12, 931–934.

2. Avsec, Z., Agarwal, V., Visentin, D., Ledsam, J.R., Grabska-Barwinska, A., Taylor, K.R., Assael, Y., Jumper, J., Kohli, P., and Kelley, D.R. (2021). Effective gene expression prediction from sequence by integrating long-range interactions. Nat Methods 18, 1196-+.

3. Tarazona, S., Arzalluz-Luque, A., and Conesa, A. (2021). Undisclosed, unmet and neglected challenges in multi-omics studies. Nat Comput Sci 1, 395–402.

4. Sammut, S.J., Crispin-Ortuzar, M., Chin, S.F., Provenzano, E., Bardwell, H.A., Ma, W.X., Cope, W., Dariush, A., Dawson, S.J., Abraham, J.E., et al. (2022). Multi-omic machine learning predictor of breast cancer therapy response. Nature 601, 623-+.

5. Cai, Z.X., Poulos, R.C., Liu, J., and Zhong, Q. (2022). Machine learning for multiomics data integration in cancer. Iscience 25.

6. Fu, Y.H., Xu, J.Y., Tang, Z.S., Wang, L., Yin, D., Fan, Y., Zhang, D.D., Deng, F., Zhang, Y.P., Zhang, H.H., et al. (2020). A gene prioritization method based on a swine multi-omics knowledgebase and a deep learning model. Commun Biol 3.

7. Swati, Z.N.K., Zhao, Q.H., Kabir, M., Ali, F., Ali, Z., Ahmed, S., and Lu, J.F. (2019). Brain tumor classification for MR images using transfer learning and fine-tuning. Comput Med Imag Grap 75, 34–46.

8. Laparra, E., Mascio, A., Velupillai, S., and Miller, T. (2021). A Review of Recent Work in Transfer Learning and Domain Adaptation for Natural Language Processing of Electronic Health Records. Yearb Med Inform 30, 239–244.

9. Kirchler, M., Konigorski, S., Norden, M., Meltendorf, C., Kloft, M., Schurmann, C., and Lippert, C. (2022). transferGWAS: GWAS of images using deep transfer learning. Bioinformatics 38, 3621–3628.

10. Cai, C., Wang, S., Xu, Y., Zhang, W., Tang, K., Ouyang, Q., Lai, L., and Pei, J. (2020). Transfer Learning for Drug Discovery. J Med Chem 63, 8683–8694.

11. Chan, H.P., Samala, R.K., Hadjiiski, L.M., and Zhou, C. (2020). Deep Learning in Medical Image Analysis. Adv Exp Med Biol 1213, 3–21.

12. Toseef, M., Olayemi Petinrin, O., Wang, F., Rahaman, S., Liu, Z., Li, X., and Wong, K.C. (2023). Deep transfer learning for clinical decision-making based on high-throughput data: comprehensive survey with benchmark results. Brief Bioinform 24.

13. Mazo, C., Bernal, J., Trujillo, M., and Alegre, E. (2018). Transfer learning for classification of cardiovascular tissues in histological images. Comput Methods Programs Biomed 165, 69–76.

14. Huang, K.Z., Xu, Z.L., King, I., Lyu, M.R., and Campbell, C. (2009). Supervised Self-taught Learning: Actively Transferring Knowledge from Unlabeled Data. Ieee Ijcnn, 481-+.

15. Ng, A. (2016). Nuts and bolts of building AI applications using Deep Learning. NIPS Keynote Talk.

16. Manolio, T.A., Collins, F.S., Cox, N.J., Goldstein, D.B., Hindorff, L.A., Hunter, D.J., McCarthy, M.I., Ramos, E.M., Cardon, L.R., Chakravarti, A., et al. (2009). Finding the missing heritability of complex diseases. Nature 461, 747–753.

17. Tam, V., Patel, N., Turcotte, M., Bosse, Y., Pare, G., and Meyre, D. (2019). Benefits and limitations of genome-wide association studies. Nat Rev Genet 20, 467–484.

18. Price, A.L., Zaitlen, N.A., Reich, D., and Patterson, N. (2010). New approaches to population stratification in genome-wide association studies. Nat Rev Genet 11, 459–463.

19. Weissbrod, O., Hormozdiari, F., Benner, C., Cui, R., Ulirsch, J., Gazal, S., Schoech, A.P., van de Geijn, B., Reshef, Y., Marquez-Luna, C., et al. (2020). Functionally informed fine-mapping and polygenic localization of complex trait heritability. Nat Genet 52, 1355–1363.

20. He, J., Wen, W., Beeghly, A., Chen, Z., Cao, C., Shu, X.O., Zheng, W., Long, Q., and Guo, X. (2022). Integrating transcription factor occupancy with transcriptome-wide association analysis identifies susceptibility genes in human cancers. Nat Commun 13, 7118.

21. He, J., Wen, W., Ping, J., Li, Q., Chen, Z., Perera, D., Shu, X., Long, J., Cai, Q., Shu, X., Zheng, W., Long, Q., Guo, X. (2023+). Transcription Factor-Linked Translocated Variants Improve Transcriptome-Wide Association Analysis for Disease Susceptibility Gene Discovery. Uneder review in Genome Medicine.

22. Wen, W., Chen, Z., Bao, J., Long, Q., Shu, X.O., Zheng, W., and Guo, X. (2021). Genetic variations of DNA bindings of FOXA1 and co-factors in breast cancer susceptibility. Nat Commun 12, 5318.

23. Wang, G., Sarkar, A., Carbonetto, P., and Stephens, M. (2020). A simple new approach to variable selection in regression, with application to genetic fine mapping. J R Stat Soc Series B Stat Methodol 82, 1273–1300.

24. Cao, C., Kwok, D., Edie, S., Li, Q., Ding, B., Kossinna, P., Campbell, S., Wu, J., Greenberg, M., and Long, Q. (2021). kTWAS: integrating kernel machine with transcriptome-wide association studies improves statistical power and reveals novel genes. Brief Bioinform 22.

25. Cowper-Sal-lari, R., Zhang, X.Y., Wright, J.B., Bailey, S.D., Cole, M.D., Eeckhoute, J., Moore, J.H., and Lupien, M. (2012). Breast cancer risk-associated SNPs modulate the affinity of chromatin for FOXA1 and alter gene expression. Nat Genet 44, 1191–1198.

26. Guo, X., Lin, W., Bao, J., Cai, Q., Pan, X., Bai, M., Yuan, Y., Shi, J., Sun, Y., Han, M.R., et al. (2018). A Comprehensive cis-eQTL Analysis Revealed Target Genes in Breast Cancer Susceptibility Loci Identified in Genome-wide Association Studies. Am J Hum Genet 102, 890–903.

27. Li, Q.Y., Seo, J.H., Stranger, B., McKenna, A., Pe’er, I., LaFramboise, T., Brown, M., Tyekucheva, S., and Freedman, M.L. (2013). Integrative eQTL-Based Analyses Reveal the Biology of Breast Cancer Risk Loci. Cell 152, 633–641.

28. Zhan, H.Y., Ahearn, T.U., Lecarpentier, J., Barnes, D., Beesley, J., Qi, G.H., Jiang, X., O’Mara, T.A., Zhao, N., Bolla, M.K., et al. (2020). Genome-wide association study identifies 32 novel breast cancer susceptibility loci from overall and subtype-specific analyses. Nat Genet 52, 572-+.

29. Cao, C., Kossinna, P., Kwok, D., Li, Q., He, J., Su, L., Guo, X., Zhang, Q., and Long, Q. (2022). Disentangling genetic feature selection and aggregation in transcriptome-wide association studies. Genetics 220.

30. Gusev, A., Ko, A., Shi, H., Bhatia, G., Chung, W., Penninx, B.W., Jansen, R., de Geus, E.J., Boomsma, D.I., Wright, F.A., et al. (2016). Integrative approaches for large-scale transcriptome-wide association studies. Nat Genet 48, 245–252.

31. Consortium, G.T. (2015). Human genomics. The Genotype-Tissue Expression (GTEx) pilot analysis: multitissue gene regulation in humans. Science 348, 648–660.

32. Pinero, J., Queralt-Rosinach, N., Bravo, A., Deu-Pons, J., Bauer-Mehren, A., Baron, M., Sanz, F., and Furlong, L.I. (2015). DisGeNET: a discovery platform for the dynamical exploration of human diseases and their genes. Database-Oxford.

33. Sondka, Z., Bamford, S., Cole, C.G., Ward, S.A., Dunham, I., and Forbes, S.A. (2018). The COSMIC Cancer Gene Census: describing genetic dysfunction across all human cancers. Nat Rev Cancer 18, 696–705.

34. Amariuta, T., Luo, Y., Gazal, S., Davenport, E.E., van de Geijn, B., Ishigaki, K., Westra, H.J., Teslovich, N., Okada, Y., Yamamoto, K., et al. (2019). IMPACT: Genomic Annotation of Cell-State-Specific Regulatory Elements Inferred from the Epigenome of Bound Transcription Factors. Am J Hum Genet 104, 879–895.

35. Crouch, D.J.M., and Bodmer, W.F. (2020). Polygenic inheritance, GWAS, polygenic risk scores, and the search for functional variants. Proc Natl Acad Sci U S A 117, 18924–18933.

36. Amariuta, T., Ishigaki, K., Sugishita, H., Ohta, T., Koido, M., Dey, K.K., Matsuda, K., Murakami, Y., Price, A.L., Kawakami, E., et al. (2020). Improving the transancestry portability of polygenic risk scores by prioritizing variants in predicted cell-type-specific regulatory elements. Nat Genet 52, 1346–1354.

37. Tong, X.Y., Xu, X.X., Huang, S.L., and Zheng, L.Z. (2021). A Mathematical Framework for Quantifying Transferability in Multi-source Transfer Learning. Advances in Neural Information Processing Systems 34 (Neurips 2021) 34.

38. Liu, T., Ortiz, J.A., Taing, L., Meyer, C.A., Lee, B., Zhang, Y., Shin, H., Wong, S.S., Ma, J., Lei, Y., et al. (2011). Cistrome: an integrative platform for transcriptional regulation studies. Genome Biol 12.

39. Stegle, O., Parts, L., Piipari, M., Winn, J., and Durbin, R. (2012). Using probabilistic estimation of expression residuals (PEER) to obtain increased power and interpretability of gene expression analyses. Nat Protoc 7, 500–507.

40. Purcell, S., Neale, B., Todd-Brown, K., Thomas, L., Ferreira, M.A., Bender, D., Maller, J., Sklar, P., de Bakker, P.I., Daly, M.J., et al. (2007). PLINK: a tool set for whole-genome association and population-based linkage analyses. Am J Hum Genet 81, 559–575.

41. Roadmap Epigenomics, C., Kundaje, A., Meuleman, W., Ernst, J., Bilenky, M., Yen, A., Heravi-Moussavi, A., Kheradpour, P., Zhang, Z., Wang, J., et al. (2015). Integrative analysis of 111 reference human epigenomes. Nature 518, 317–330.

42. Ernst, J., and Kellis, M. (2012). ChromHMM: automating chromatin-state discovery and characterization. Nat Methods 9, 215–216.

43. Alzubaidi, L., Zhang, J., Humaidi, A.J., Al-Dujaili, A., Duan, Y., Al-Shamma, O., Santamaría, J., Fadhel, M.A., Al-Amidie, M., and Farhan, L. (2021). Review of deep learning: Concepts, CNN architectures, challenges, applications, future directions. Journal of Big Data 8, 1–74.

44. Hayden, M.S., and Ghosh, S. (2008). Shared principles in NF-kappaB signaling. Cell 132, 344–362.

